# Structure of the Fanconi Anemia Core–UBE2T complex poised to ubiquitinate bound FANCI–FANCD2

**DOI:** 10.1101/854158

**Authors:** Shengliu Wang, Renjing Wang, Christopher Peralta, Ayat Yaseen, Nikola P. Pavletich

## Abstract

The Fanconi Anemia (FA) pathway is essential for the repair of DNA interstrand crosslinks (ICLs). The pathway is activated when a replication fork stalls because of an ICL or other replication stress. A central event in pathway activation is the mono-ubiquitination of the FANCI-FANCD2 (ID) complex by the FA Core complex, a ubiquitin ligase of nine subunits. Here we describe the cryo-EM structures of the 1.1 MDa FA Core at 3.1 angstroms, except for the FANCA subunit at 3.4, and of the complex containing Core, ID and the UBE2T ubiquitin conjugating enzyme at 4.2 angstroms. The Core has unusual stoichiometry with two copies of FANCB, FAAP100, FANCA, FAAP20, FANCG, FANCL, but only a single copy of FANCC, FANCE and FANCF. This is due to homodimers of FANCA and FANCB having incompatible 2-fold symmetry, resulting in an overall asymmetric assembly of the other subunits. The asymmetry is crucial, as it prevents the binding of a second FANC-C-E-F sub-complex that inhibits UBE2T recruitment by FANCL, and instead creates an ID binding site. The single active FANCL-UBE2T binds next to the FANCD2 ubiquitination site, prying open the FANCI-FANCD2 interface within which the ubiquitination sites are buried. These structures and biochemical data indicate a single active site ubiquitinates FANCD2 and FANCI sequentially, shedding light on a central event in the FA pathway.

---

Fanconi Anemia (FA) is a recessive syndrome of chromosome instability and cancer predisposition^1^. FA cells are hypersensitive to DNA inter-strand crosslinking agents (ICLs), and the 22 FA complementation group proteins define a pathway essential for ICL repair^2, 3^. The FA pathway is activated by replication forks that stall at an ICL, and it acts in concert with other repair pathways to excise the ICL from one DNA strand, bypass it from the other and repair the lesion by homologous recombination^2, 4^. FA proteins are also implicated in the protection and recovery of replication forks that stall at common fragile sites or during other forms of replication stress^1, 2, 5, 6^. A key event in pathway activation is the mono-ubiquitination of the FANCI-FANCD2 complex (thereafter ID complex) by a nine-subunit ubiquitin ligase named FA Core complex ^2, 7^. Mono-ubiquitination converts the ID complex, which has DNA binding activity^8–10^, from an open trough-like shape to a closed ring that encircles the DNA^11^.

The Core complex consists of FANC-A, B, C, E, F, G, and L, and the non-FA proteins FAAP100 and FAAP20^2^. FANCL contains a RING domain that recruits the UBE2T (FANCT) ubiquitin conjugating E2 enzyme^12, 13^ as with other RING E3 enzymes^14^. The roles of the other Core complex subunits are poorly understood. Except for FANCB and FAAP100, they are predicted to consist of helical repeats as demonstrated by the crystal structures of FANCE and FANCF C-terminal domains ^15–17^. Biochemical and genetic studies have identified a number of subunit interactions, including between FANCE and FANCD2^18^, and subcomplexes consisting of FANC-A-G-FAAP20, FANC-C-E-F, and FANC-B-L-FAAP100 ^19–26^. A 2D EM analysis of the FANC-B-L-FAAP100 sub-complex, which can support some level of mono-ubiquitination in vitro, showed a 2-fold symmetric dimer of hetero-trimers, and suggested that it assembles with two copies of FANC-C-E-F for the symmetric mono-ubiquitination of FANCI and FANCD2^27^.

However, these studies have not shed light upon the mechanism of ID ubiquitination, including the question of how the Core complex accesses the ubiquitination sites that are sequestered inside the FANCI-FANCD2 interface^10, 11^. To address these questions, we have determined the cryo-EM structure of the 9 subunit Core complex, as well as the structures of the Core-ID and Core-ID-UBE2T complexes.

## Structure determination

The human FA Core complex ubiquitin ligase was produced from a stably transfected HEK-293F cell line overexpressing eight subunits (FANC A, B, C, E, F, G, L, FAAP100) (Extended Data Fig. 1a). The purified Core complex also contains the endogenous FAAP20 subunit, as revealed by the cryo-EM density and subsequently confirmed by immunoblotting (Extended Data Figs 1b to c). Cryo-EM data was collected at a Core complex concentration of 1 to 1.3 *μ*M. The initial consensus reconstruction of 671,972 particles with the program RELION3 extended to an overall resolution of 3.4 Å as determined by the gold-standard fourier shell correlation (FSC) procedure (Extended Data Fig. 2a; Extended Data Table 1). Focused 3D refinements using seven masks yielded reconstructions with the highest common resolution of 3.1 Å, except for portions of the FANCA dimer, which has at least two conformations (Extended Data Figs 2a and b). For the most rigid FANCA conformation (309,725 particles) the reconstruction extended to 3.4 Å and 3.7 Å for the two FANCA copies in the complex (Extended Data Figs 2a and b; cryo-EM density for subunit interfaces is shown in Extended Data Figs 3a to k).

**Figure 1.**
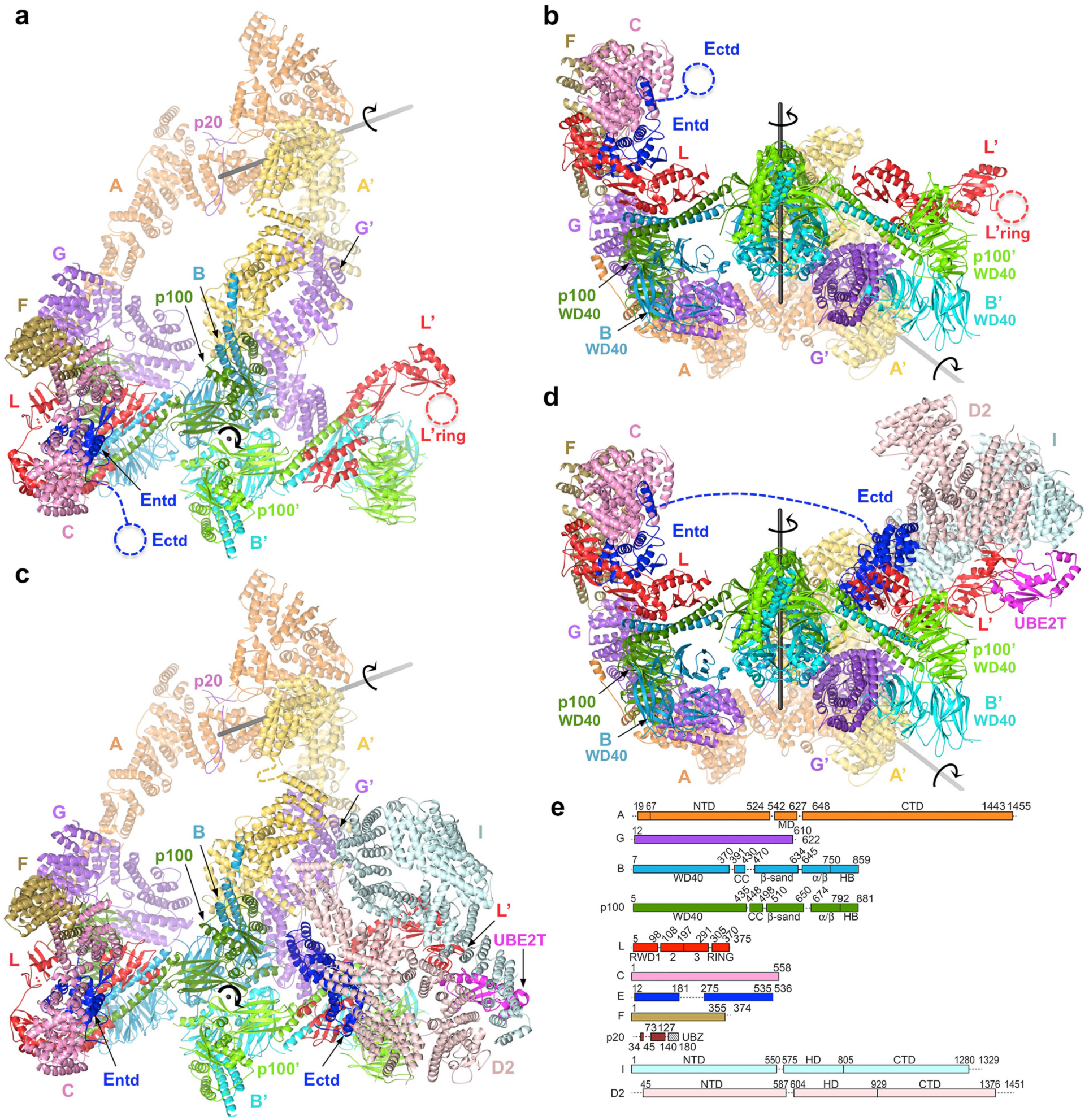
Overall Structures of the Core and Core-UBE2T-ID complexes. **a,** Core complex cartoon representation with individual subunits (labeled) colored according to schematic in (d), except for the homodimers of FANCA and FANCB-FAAP100 where the second protomer of each is colored in lighter shade. Dashed circles indicate disordered domains. Where applicable, the second copy of a subunit is indicated by the symbol prime on the label. View looking down the 2-fold symmetry axis (marked) of the FANCB-FAAP100 hub. The FANCA dimer axis is shown as a gray line. **b,** View looking up the vertical axis of a. **c,** Cartoon representation of the Core-UBE2T-ID complex in the same orientation as a. FANCI is light cyan, FANCD2 light pink, and UBE2T magenta. View is looking into the ID trough. Dashed blue line indicates the disordered residues between the FANCE NTD and CTD domains. **d,** View looking up the vertical axis of c. e, Linear representation of Core complex subunits. Dashed lines indicate disordered segments that are longer than 10 residues. Unnamed rectangles consist mostly of helical repeats.

**Figure 2.**
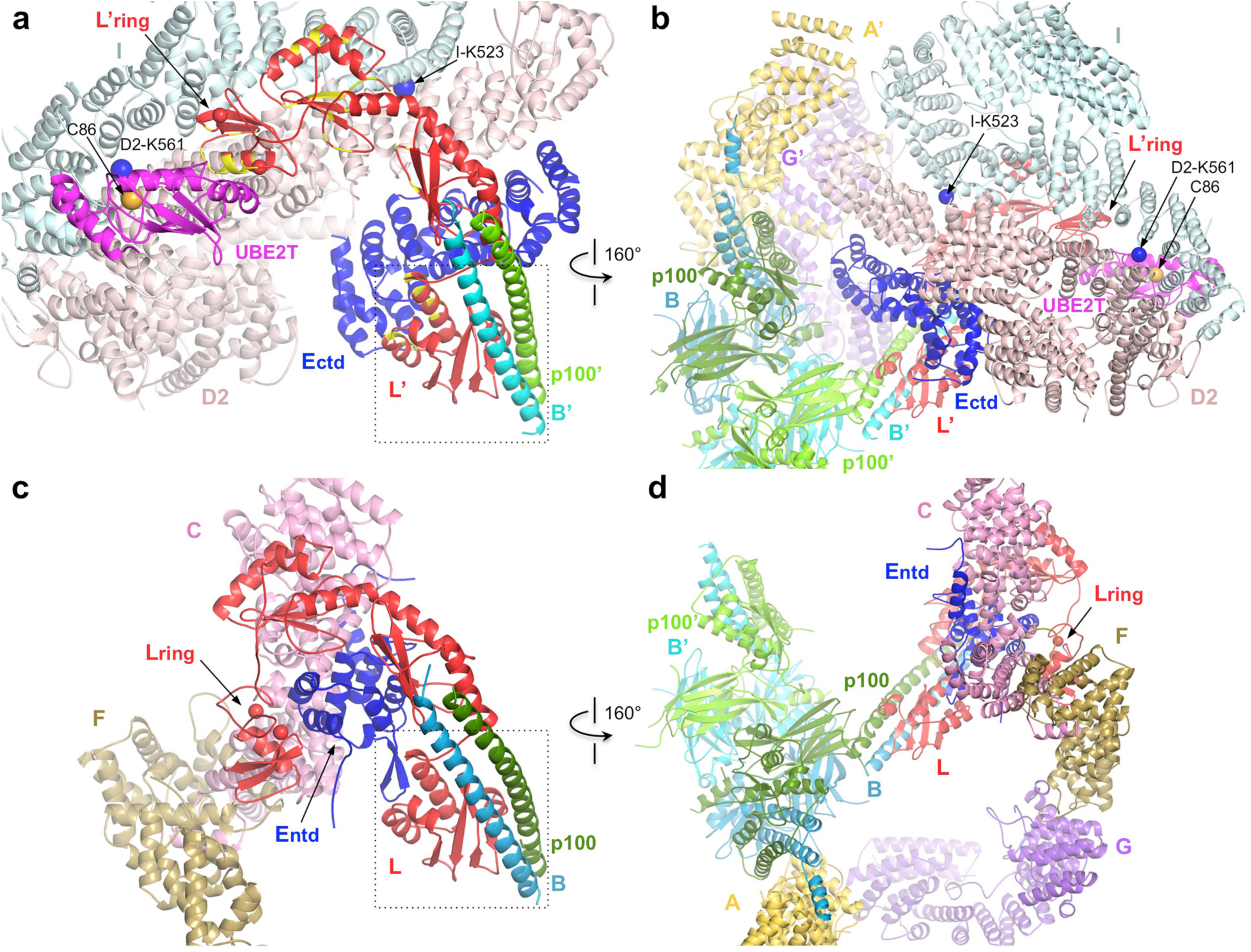
Comparison of the active and inactive sides of the pseudo-dimeric Core-UBE2T-ID complex. **a,** Close-up view of ID on Core. For clarity only subunits or portions thereof in the immediate viccinity of FANCL’ are shown. FANCL’ residues with yellow secondary structure correspond to residues that are sequestered within 3.8 Å of the FANC-C-ENTD-F sub-complex of the inactive side shown in (c), aligned on the FANC-L-B-FAAP100 residues enclosed in the dotted box. Ubiquitination sites (FANCI Lys523, FANCD2 Lys561) and UBE2T active site Cys86 are indicated with spheres in blue and orange, respectively. View looking from bellow the plane of Figure 1c (at the exterior bottom of the ID trough). **b,** View of Core-UBE2T-ID in an orientation similar to figure 1c, also showing the proximity of FANCA’ to FANCI. **c-d,** The inactive side of the complex aligned on the active side views of (a) and (b) as described in (a).

The Core-ID data was collected from samples that contained 0.8 to 1.2 *μ*M Core, 1.5 to 3 *μ*M ID complex and 3 *μ*M of 58 base pair nicked DNA. Although DNA was shown to promote ID ubiquitination^21, 28, 29^, there is no DNA density on the ID complex. The Core-ID-UBE2T samples additionally contained 6 *μ*M UBE2T harboring the C86K active site mutation through which it was conjugated to ubiquitin. The maps show no ubiquitin density and we presume it is disordered. Approximately one third of Core complex particles in each data set contained the ID complex. The structures were determined with a focused reconstruction approach similar to the isolated Core complex, with a highest common resolution of 4.2 Å from 114,249 Core-ID-UBE2T particles and 4.5 Å from 76,111 Core-ID particles (Extended Data Figs 2c and d; Extended Data Table 1).

## Overall Structures of the FA Core and FA Core-UBE2T-ID complexes

The Core complex has a flat appearance with approximately 315 Å and 240 Å in two dimensions, but only 165 Å in the third. The bulk of its mass is in a relatively compact body from which three arms made up of two FANCA molecules extend and meet at a distal tip, creating a cage-like structure (Figs 1a and b).

The complex has an unusual stoichiometry with two copies of FANCB, FAAP100, FANCA, FANCG, FANCL, but only a single copy of FANCC, FANCE, FANCF (Figs 1a and b). The body of the particle is organized around a dimer of FANCB-FAAP100 heterodimers. Their C-terminal portions form a locally 2-fold symmetric center, while their N-terminal portions extend out to opposite sides in non-symmetric arrangements. At one side (left on Fig. 1a), the FANCB-FAAP100 N-terminal portions assemble with the FANC-G-L-C-E-F proteins, but at the other side only with FANC-G-L (thereafter these subunits designated by the prime symbol).

Besides the lack of FANC-C-E-F, this side also differs in the conformation of FANCL’ as well as the position, orientation and packing contacts of FANCG’ (Figs 1a and b). This impacts the disposition of the two FANCA molecules that extend from the body. FANCA is the only other subunit that forms a homodimer, but its symmetry is unrelated to that of FANCB-FAAP100, as the two 2-fold symmetry axes are 131 Å apart and at a 55° angle (Figs 1a and b). In addition, the two FANCA protomers have different conformations associated with the differentially positioned FANCG and FANCG’ on which they anchor.

The asymmetric composition and arrangement of the Core subunits are essential for binding to the ID complex, which occupies the overall area where a second FANC-C-E-F would have been in a symmetric complex (Figs 1c and d). Central to ID binding is FANCL’. It interacts extensively with FANCI and to a lesser extent with FANCD2, and it also binds to the FANCE C-terminal domain (FANCE^CTD^), which in turn interacts with FANCD2 as well^18^ (Figs 2a and b). FANCE^CTD^ is connected to the FANCE N-terminal domain (FANCE^NTD^) at the other side of the body through a 92 amino acid unstructured segment spanning 110 Å, and it is disordered in the apo-Core complex (Figs 1a and b). FANCL also recruits the UBE2T enzyme, juxtaposing its active site with the FANCD2 ubiquitination site (Fig. 2a). None of these FANCL’ interactions can be made by FANCL because it is sequestered by the FANC-C-E^NTD^-F sub-complex (compare Figs 2a and b to c and d). To reflect this, we will be referring to the FANCL-containing side of the body as inactive, and that with FANCL’ active.

## Core complex assembly

The most extensive intermolecular protein-protein interface is between FANCB and FAAP100, which intertwine to bury ∼15,400 Å^2^ of total surface area (secondary structures and intermolecular contacts are summarized in Extended Data Figs 4a to h). The structures of the two proteins are built from the same five domains and likely diverged from a common ancestor. Each starts with an N-terminal WD40 domain that packs with the paralog’s WD40 domain, followed by long helix that forms an intermolecular coiled-coil, then a globular domain consisting of a β sandwich, an α*/*β domain, and a helical bundle (HB) (Fig. 1e and Extended Data Figs 5a to b). Each of these structural domains contributes to FANCB-FAAP100 association. The coiled-coil and β sandwich domains interact with their paralogous mates with an approximate pseudo-2-fold symmetry, but the WD40-WD40 and the α*/*β−α*/*β packing arrangements are serendipitous (Extended Data Figs 5a to b). The two proteins have also accumulated insertions such as a FANCB helical bundle insertion that interacts with FANCA.

The FANCB-FAAP100 heterodimer forms a homodimer as reported^19, 27^. Homodimerization is through the packing of the β sandwich and α*/*β domains of each protein with the corresponding domain of the second protomer (Extended Data Fig. 6a). This interface is smaller than that of hetero-dimerization, burying ∼4,500 Å^2^. The β sandwich and α*/*β domains at the globular center obey proper 2-fold symmetry, but the WD40-WD40 pairs at the periphery end up rotated by ∼50° and shifted by ∼25 Å (Extended Data Figs 6b and c). The WD40-WD40 pairs do not pack with the globular center, and their positions are influenced by other core subunits.

This dimer of hetero-dimers then serves as an assembly hub, with the two WD40-WD40 pairs likely flexibly attached to the hub or sampling multiple conformations (schematic of assembly sequence in Extended Data Fig. 7). FANCL binds readily to this hub as reported^19, 27^, and in fact on the active side of the Core complex, FANCL’ contacts no subunits other than FANCB-FAAP100 (Figs 1a and b). FANCL, which consists of three RWD domains followed by the RING domain^12^, uses its RWD1 and RWD2 domains to bind to the FANCB-FAAP100 coiled coil, burying ∼4,300 Å^2^ (Figs 2a and c; Extended Data Figs 8a to g).

The next steps of the assembly likely involve the pre-assembled FANC-C-E-F and FANC-G-A-FAAP20 sub-complexes^19, 20, 22, 24^. The FANC-C-E-F subunits, which consist of helical repeats of the HEAT family, come together through large, intertwined interfaces, with the central FANCC binding to both FANCF and FANCE^NTD^ and burying ∼6300 and ∼5800 Å^2^ of surface area, respectively (Extended Data Figs 9a to e).

The FANCA-FANCG sub-complex forms through a highly conserved N-terminal FANCA segment (residues 19 to 67) binding to FANCG as predicted by yeast-2-hybrid^24^ (3,850 Å^2^ buried; Extended Data Figs 10a to f). This segment and its associated FANCG is rotated by ∼132° relative to the rest of FANCA in the two sides of the Core complex, and appears to be flexibly linked (Extended Data Fig. 11). After this segment, FANCA contains three structural domains with intervening linkers unstructured (Fig. 1e). The larger NTD and CTD, which are made up of HEAT repeats, pack together in FANCA, with the smaller helical MD packing with the CTD (Extended Data Fig. 11). By contrast, the NTD and CTD are uncoupled in the active-side FANCA’, and the MD is disordered. FANCG consists of TPR repeats and has a structural zinc atom.

The structure suggests that the FANC-C-E-F and FANC-A-G subcomplexes cooperate in binding to the FANC-B-P100-L hub (step 2 in Extended Data Fig. 7). On the inactive side, FANCG packs with the FAAP100 WD40 domain (3,900 Å^2^ buried; Extended Data Fig. 10a and i). We presume this interaction is not stable enough to persist by itself, because the active-side FAAP100’ WD40 domain is devoid of FANCG even though it is fully accessible. Similarly, the binding of the FANC-C-E-F sub-complex to the hub appears not be stable enough to persist by itself. Its contacts, which are made by FANCC and FANCE (2,500 and 2,650 A^2^, respectively) to all four domains of FANCL (RWD 1 to 3, and RING; Extended Data Fig. 8) can in principle also be made to the fully accessible active side FANCL’, which nonetheless is devoid of FANC-C-E-F. What likely stabilizes the transient association of FANC-A-G and FANC-C-E-F with the FANCB-FAAP100-FANCL hub is avidity, through an interaction between FANCG and two highly conserved FANCF loops whose mutation was shown to compromise Core complex stability and FANCD2 ubiquitination^15^ (Extended Data Fig. 10a and g).

Once a FANCG molecule is anchored on the FANC-B-P100-L hub, its associated FANCA dimer brings in FANCG’ at a high local concentration (Extended Data Fig. 7). However, the FANCG’ cannot be engaged by the emerging Core complex in a 2-fold symmetric manner because its binding site would be at least 100 Å away, at a different end of the hub (Extended Data Fig. 12). And, while dissociation of the FANCA dimer would make this possible, this is unlikely as the FANCA-FANCA dimerization interface contains an FA mutation, indicating an essential role for dimerization (Extended Data Fig.13).

Rather, the second FANCG’ that comes along binds to an entirely different site on the hub, making a new set of multivalent interactions. By contrast to FANCG that packs with the FAAP100 WD40 domain, FANCG’ contacts the FANCB’ WD40 and the FANCB α*/*β domains, burying 1,250 and 840 A^2^, respectively (Extended Data Fig. 10b). This relatively smaller interface is augmented by the FANCG’-associated FANCA’ contacting the HB domains of FANC-B and FAAP100, burying 2,800 Å^2^ in an interface not possible with the inactive side FANCA (Extended Data Fig. 11d). Also not possible with FANCA is the packing of the FANCA’ C-terminus with the FANCG’ C-terminus (Extended Data Fig. 10h). This results in the closing of a FANCA-FANCG’ solenoid circle, and the uncoupling of the FANCA’ NTD-CTD interface.

Crucially, the switch from FANCG packing with FAAP100 WD40 at the inactive side to FANCG’ packing with FANCB’ WD40 is associated with, and likely causes, the different FANCB’-FAAP100’ WD40 domain positions. The functional consequence of this is that the active side of the complex cannot form the bridge contacts between FANCG and the FANC-C-E-F sub-complex. Without the avidity from this interaction, FANC-C-E-F cannot stably bind to and sequester FANCL’ at the active side of the Core complex (Extended Data Fig. 7).

This assembly sequence makes the prediction that if fully 2-fold symmetric body particles exist, they should consist of two copies of FANC-B-P100-L-C-E-F-G, with the FANCA dimers flexibly tethered. Indeed, using successive 3D classifications we identified 7,331 particles (∼1 % of dataset) whose 3D reconstruction in point group C1 is fully symmetric and shows clear density for two copies of FANC-B-P100-L-C-E-F-G with only residual FANCA^NTD^ density (6.4 to 7.6 Å; Extended Data Fig. 14). We presume that this is a dead-end side reaction resulting from the binding of a second set of FANC-A-G and FANC-C-E-F sub-complexes from solution before the intramolecular association of FANCG’ with the hub is completed.

## ID and UBE2T binding

The paralogous FANCI and FANCD2 each consist of a long helical-repeat solenoid (henceforth NTD), followed by a helical domain (HD) that reverses the direction of the solenoid, and a C-terminal helical-repeat domain (CTD; Fig 1e). The two proteins come together along their NTD solenoids in an antiparallel manner to form an open trough-like structure, as reported for the isolated mouse and human ID structures^10, 11^.

The Core complex has two major recruitment points for the ID complex. In keeping with biochemical reports^18^, the FANCE^CTD^ binds to the N-terminal portion FANCD2^NTD^, interacting with the exterior bottom of the trough-like structure (Figs 3a and b). It uses most of its seven heat repeats, except for a gap near the center, to contact six FANCD2 repeats, burying ∼2,120 Å^2^ (Fig. 3b). The other is FANCL’, which encircles the ID-FANCE^CTD^ complex partway around the bottom of the trough (Fig. 3a). The FANCL’ RWD1 domain binds to the FANCE^CTD^, and after RWD2 crosses the I-D interface, the RWD3 and RING domains run along the FANCI^NTD^ staying close to the I-D interface.

**Figure 3.**
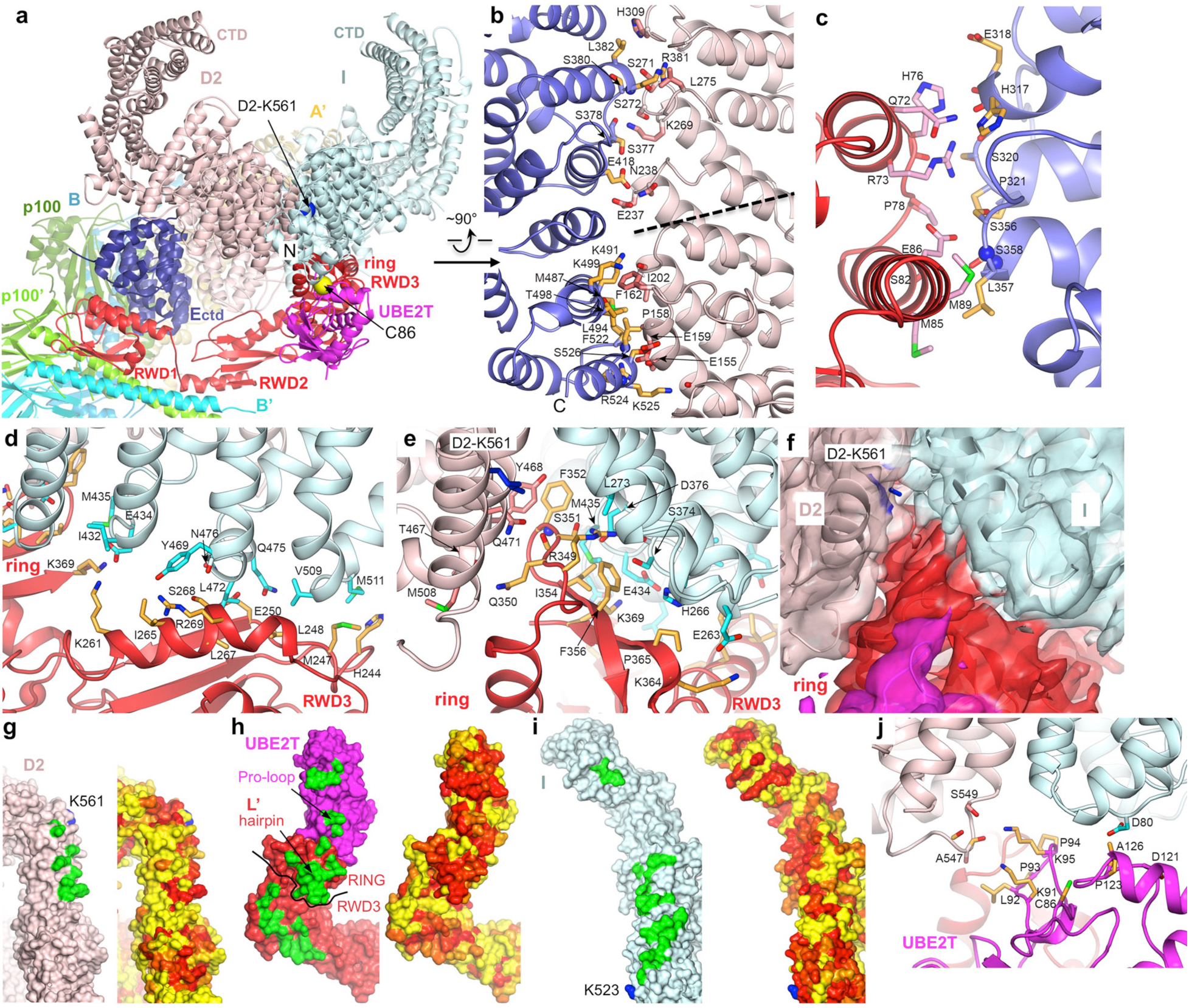
ID binding to Core-UBE2T. **a,** View of FANCECTD-FANCL’-UBE2T partway encircling the exterior bottom of the trough-like ID structure. View looking down the end of the trough-like shape that has the FANCI N-terminus (labeled). **b,** FANCD2-FANCECTD interface showing side chain and backbone groups within 3.8 Å of the binding partner (side chains but not backbone groups are labeled). Dashed line indicates the hinge region about which the FANCD2 N-terminal segment rotates away from FANCI. **c,** Side chain and backbone groups at the interface between the FANCL’ RWD1 domain and the FANCECTD. **d,** Residues at the interface between FANCI and the FANCL’ RWD3 domain. Residues at the RING interface are not labeled as they are shown in the next panel (e). **e,** Side chain and backbone groups at the interface between ID and the RING domain of FANCL. **f,** 4.2 Å cryo-EM density (semi-transparent) in an orientation similar to e and colored according to distance from each protein. Most of UBE2T (magenta) is clipped for clarity. **g to i,** Molecular surface representations of FANCD2, FANCL’-UBE2T and FANCI viewed from the perspective of the interacting partner (FANCL’-UBE2T perspective for FANCI and FANCD2, and ID perspective for FANCL’-UBE2T), marking intermolecular contacts green in left panels (FANCL’UBE2T contacts for FANCI and FANCD2, and ID contacts for FANCL’-UBE2T), and colored by conservation (yellow to red for low to high conservation) in right panels. Select elements of FANCL’UBE2T are labeled. Dark curve on FANCL’ delineates its RWD3 and RING domains. j, Side chain and backbone groups at the interface between ID and the UBE2T proline-rich loop.

The interface between the FANCL’ RWD1 domain and FANCE^CTD^ is relatively smaller (∼900 Å^2^) though it has a high density of contacts, including between five FANCE^CTD^ backbone groups and FANCL’ side chains in a mostly buried environment (Fig. 3c). Nevertheless, this interface does not suffice for the FANCE^CTD^ to bind to FANCL’ in the absence of the avidity provided by ID-Core interactions.

The FANCL’ RWD3 binds exclusively to FANCI (∼1,150 Å^2^ buried; Fig. 3d), while the RING domain interacts with both FANCI and FANCD2 (∼1,360 Å^2^ buried; Fig. 3e). Key to RING domain binding is a hydrophobic hairpin that inserts deep into the I-D interface coming from the bottom of the trough, in the vicinity of the FANCD2 Lys561 ubiquitination site embedded inside the ID interface (Fig. 3f). The hairpin Phe352, Ile354 and Phe356 side chains pack with hydrophobic residues from both FANCI and FANCD2, while Arg349 and Ser351 are within hydrogen-bonding distance of polar side chain and backbone groups (Figs 3e and f). While the residues at all the FANCL’-ID interfaces are overall well conserved, those at the RING interface exhibit the highest level of conservation, indicating it has a central role in ID ubiquitination (Figs 3g to i).

UBE2T binds to the FANCL’ RING domain in an orientation very similar to the isolated RING domain-UBE2T crystal structure^13^. This results in ID contacts from two UBE2T patches flanking the active site Cys86, which is ∼24 Å away from the FANCD2 Lys561 (Fig. 3a). Compared to FANCL-ID contacts, those of the UBE2T-ID interface are relatively minor (∼410 Å^2^ buried; Fig. 3j). Nevertheless, there are parallels with the RING hairpin in that UBE2T has a proline-rich loop (residues 92 to 95) that reaches partway into the ID interface, though not as deep as the RING hairpin. This loop is an insertion to the E2 fold unique to UBE2T, though a few other E2s contain distinct insertions at the equivalent location.

## ID conformation

On binding to the Core complex, the I-D interface opens up through a relative rotation of FANCI and FANCD2 by ∼8° about an axis running along their NTD-NTD interface (Fig. 4a; Extended Data Fig. 14). This is coupled to and likely caused by the FANCL’ RING hairpin inserting into the I-D interface, which cannot accommodate the hairpin in the free ID conformation (Fig. 4b).

**Figure 4.**
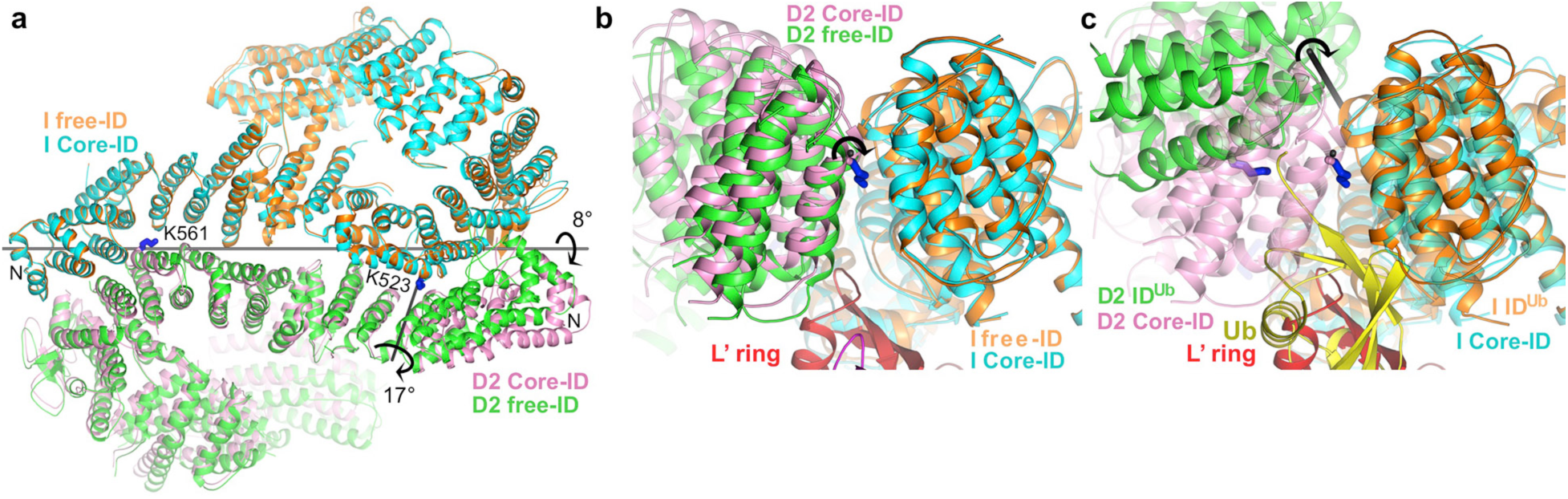
ID conformational changes on Core complex binding. **a,** Superposition of Core-bound ID and free ID aligned on their respective FANCI NTD domains, colored as marked. The horizontal gray line indicates the axis of rotation about which the FANCINTD-FANCD2NTD interface opens up on Core binding, while the shorter near vertical line marks the rotation axis of the FANCD2 N-terminal residues (45 to 205), as they get peeled away from FANCI by FANCECTD. **b,** Superposition viewed down the horizontal rotation axis of a, which now is perpendicular to the figure plane (marked with circular arrow), showing that the FANCL’ RING hairpin cannot fit into the interface in the free ID conformation. **c,** Superposition of Core-bound ID and the ubiquitinated IDUb aligned on the FANCINTD. For clarity, Core-bound ID is semi-transparent. The ubiquitin attached to FANCD2Ub (yellow) overlaps with the FANCL’ RING in this superposition. The rotation axis through which the Core-bound I-D interface further opens is shown as a gray stick.

A second substantial change is the conformation of the N-terminal ∼160 residue FANCD2 segment, which rotates by ∼17° away from FANCI (Fig. 4a). This is likely a direct consequence of binding to FANCE^CTD^, whose contacts span the FANCD2 rotation hinge (Fig. 3b). As the FANCE^CTD^ structure is essentially unchanged from the crystal structure of the isolated domain^18^, it acts as a rigid scaffold to remodel the region of FANCD2. The functional consequence of this change is a substantial loss of I-D interface contacts, with the surface area buried by this N-terminal segment reduced from ∼1,290 Å^2^ in free ID to 360 Å^2^. Together with the opening up of the rest of the NTD-NTD interface, these conformational changes reduce the total buried area at the I-D interface by ∼28% (4,930 versus 3,550 Å^2^).

The opening of the NTD-NTD interface changes the conformation of the ID complex partway towards the mono-ubiquitinated ID product (henceforth ID^Ub^), whose interface is opened up along a similar rotation axis to accommodate the ubiquitin moieties inside the NTD-NTD interface^11^ (Fig 4c). Thus, while the ID^Ub^ interface has to open up by 59° relative to free ID, this rotation is reduced to 50° relative to Core-bound ID.

## Insights into ID mono-ubiquitination

Previous findings on the dimerization of the FANCB-FAAP100-FANCL sub-complex had suggested that the Core complex mono-ubiquitinates FANCI and FANCD2 simultaneously^27^. However, our work shows that one of the two FANCL subunits is sequestered by FANC-C-E-F, indicating that a single FANCL mono-ubiquitinates FANCD2 and FANCI sequentially. The proximity of UBE2T Cys86 to FANCD2 Lys561, coupled to the clear line of sight between the two, suggests that FANCD2 mono-ubiquitination happens first. Accordingly, at the earliest time points of a reaction time course only FANCD2 shows detectable mono-ubiquitination, with ubiquitinated FANCI appearing after a substantial fraction of ubiquitinated FANCD2 has accumulated (compare 2.5 and 20 min in Fig. 5a). This is not due to FANCI mono-ubiquitination having an intrinsically slower catalytic rate, as free FANCI gets ubiquitinated with kinetics very similar to FANCD2 of the ID complex (Fig. 5b, third panel). In fact, FANCI ubiquitination exhibits similarly rapid kinetics when assembled with mono-ubiquitinated FANCD2 (ID^-/Ub^), confirming that FANCD2 ubiquitination is a pre-requisite for FANCI ubiquitination in the ID complex (Fig. 5b, first panel). As reported, free FANCD2 mono-ubiquitination is exceedingly slow (Fig. 5b, fourth panel), presumably because the FANCD2-FANCE^CTD^ fails to dock on FANCL’ in the absence of FANCI.

**Figure 5.**
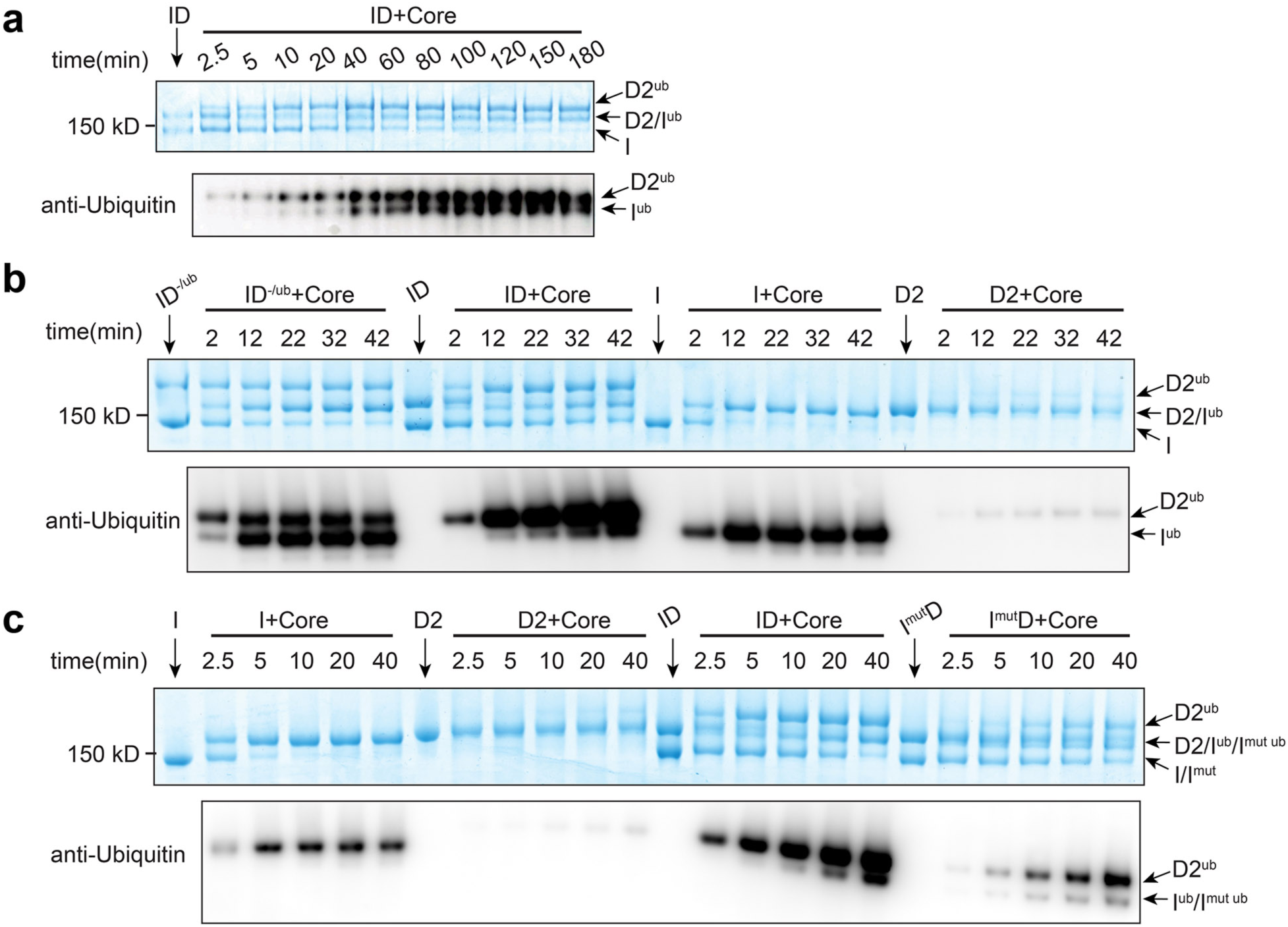
ANCD2 and FANCI are ubiquitinated sequentially. **a**, Ubiquitination time course of ID (1 *μ*M) by recombinant Core complex (0.1 *μ*M) and UBE2T (1.5 *μ*M). The positions of substrates and products are marked. **b**, Reaction time course comparing the ID complex, a complex assembled with ubiquitinated FANCD2 and non-ubiquitinated FANCI (ID^-/Ub^), and the individual proteins (all at 1 *μ*M). **c**, The R1285Q FA mutation in FANCI (FANCI^mut^) reduces ubiquitination of both FANCI and FANCD2 in the ID complex comparably but has no effect on free FANCI (all at 1 *μ*M). **a-c**, Reactions were run on the same SDS-PAGE gel for direct comparison. They were visualized by Coomassie staining (top panel) or immunoblotting with a ubiquitin-specific antibody (bottom panel). **a-c**, The experiments were repeated three times.

While the ID NTD-NTD interface is partially opened up by Core complex binding, FANCD2 ubiquitination would require further opening to accommodate the ubiquitin and also to allow UBE2T to pivot closer to Lys561. In principle, the ubiquitin attached to UBE2T may fulfill this function. Although our cryo-EM samples contained the UBE2T C86K mutant that was stably attached to ubiquitin through an isopeptide bond, an approach successfully utilized for the UbcH5A E2^30^, it is possible that this mutant does not recapitulate all conformational aspects of UBE2T.

Another possibility is that the weakened I-D interface can further open transiently to allowing Lys561 ubiquitination, but such intermediates may be too few and heterogeneous to identify in our cryo-EM samples. In the structure of ID^Ub^, the complete opening of the NTD-NTD interface is associated with the juxtaposition of the FANCI and FANCD2 CTD domains, which form a new I-D interface stabilized by an intermolecular β sheet zipper of previously unstructured C-terminal extensions^11^. The FANCI extension contains an FA mutation (R1285Q)^29, 31^ that the ID^Ub^ structure suggests would destabilize the zipper^11^. This mutation also reduces levels of ubiquitinated FANCD2 in cells^31^, suggesting that the formation of the β sheet zipper is an intermediate in FANCD2 ubiquitination. Since the mutant ID^Ub^ may also be more susceptible to de-ubiquitination by USP1 in cells, we assayed the I^R1285Q^D complex and found that the mutation reduces ubiquitination of both FANCD2 and FANCI substantially and comparably, whereas it has no effect on the ubiquitination of the isolated FANCI (Fig. 5c). This suggests that the FANCI extension may function to transiently capture the FANCD2 CTD, which exhibits substantial mobility both in the Core-bound and free^11^ ID complexes, resulting in a product-like opening of the NTD-NTD interface.

Once FANCD2 is mono-ubiquitinated and the ubiquitin packs with the FANCI^NTD11^, the FANCL RING would dissociate due to steric clashes with the ubiquitin (Fig. 4c). The FANCD2 ubiquitin would also act as a wedge that prevents the NTD-NTD interface from closing, thus exposing the FANCI Lys523 ubiquitination site. We do not yet know the mechanism of FANCI mono-ubiquitination. However, the similarly fast ubiquitination kinetics of ID^-/Ub^ and free FANCI indicate that it is independent of FANCD2-FANCE^CTD^ (Fig. 5b). In the ID^Ub^ complex, the opening of the interface drastically changes the relative positions of FANCI^NTD^ and FANCD2^NTD^, making their simultaneous interaction with FANCL^RWD3^ and FANCE^CTD^, respectively, no longer possible, suggesting a mechanism for the release of the ID^Ub^ product.

## Conclusion

The structures show that the Core complex is an asymmetric assembly, and explain how asymmetry can arise from nominally homodimeric subunits. The asymmetry is central to Core function, as it prevents the binding of a second FANC-C-E-F sub-complex that has a hitherto unknown inhibitory activity, and it also creates the proper ID binding site. The structures also show how a ubiquitin ligase can remodel its substrate to access the ubiquitination site.

## Acknowledgments

We thank the staff of the MSKCC Cryo-EM facility, the NYSBC Simons Electron Microscopy Center, and the HHMI Cryo-EM facility for help with data collection. Supported by HHMI and National Institutes of Health grant CA008748.

## Author Contributions

S.W. carried out the biochemical experiments and collected and analyzed the cryo-EM data,

R.W. prepared the ID complex, C.P. and A.P. did cell culture and protein purification, and N.P.P. analyzed the data and wrote the manuscript.

## Author Information

The authors declare no competing financial interests. Correspondence and requests for materials should be addressed to N.P.P.

## METHODS

### Protein Expression and purification

Full length human FANCA, FANCB, FANCC, FANCE, FANCF, FANCG, FANCL and FAAP100 were inserted into three modified pCDNA3.1 plasmids with various drug resistance genes that were designed for multi-gene expression via homing endonuclease (HE) site. All contain an N-terminal Flag tag except for FANCA that has a C-terminal Flag tag. The three expression vectors containing FANCA-FANCG, FANCB-FANCL-FAAP100 and FANCC-FANCE-FANCF were transfected into HEK 293F cells and selected by drugs for 30 days. A cell line with stably and highly expression of all eight subunits was then adapted for suspension culture. Cells were lysed in 50 mM Tris-HCl, 150 mM NaCl, 10% (v/v) glycerol, 1 mM DTT, pH 8.0 and protease inhibitors. After Flag-affinity, anion exchange and size-exclusion chromatography, the FA core complex was concentrated by ultrafiltration (Amicon) to approximately 10 mg/ml (9 µM) in 20 mM Bicine, 150 mM NaCl, 1mM DTT, pH 8.0 buffer and stored in small aliquots at −80 °C.

Human FANCI and human FANCD2 were expressed and purified as described^10, 11^. The R1285Q mutant of human FANCI (FANCI^mut^) was generated by using QuickChange Lightning Site-Directed Mutagenesis Kit (Agilent), and was cloned with a N-terminal cleavable Flag tag into the pCDNA 3.1 vector. FANCI^mut^ was transiently transfected into HEK293 cells cultured in suspension for 2 days. Cells were lysed in 50 mM Tris-HCl, 300 mM NaCl, 10% (v/v) glycerol, 1 mM DTT, pH 8.0 and protease inhibitors. After Flag pull-down and Flag tag cleavage, FANCI^mut^ was mixed with purified FANCD2 at one molar ratio, followed by concentration and exchange into buffer containing 150 mM NaCl. The complex was further purified by gel-filtration chromatography and concentrated by ultrafiltration (Amicon) to 11 µM in 20 mM Tris-HCl, 150 mM NaCl, 10% (v/v) glycerol, 1 mM DTT, pH 8.0 and stored in small aliquots in −80 °C.

Human UBE2T gene was cloned into a pGEX 6p-1 vector with N terminal GST tag followed by a HRV 3c site. The UBE2T C86K mutant (UBE2T^C86K^) was generated using QuickChange Lightning Site-Directed Mutagenesis Kit (Agilent). UBE2T^C86K^ was overexpressed in BL21 (DE3). It was purified using GST affinity chromatography followed by GST tag cleavage, cation exchange (Hitrap SP column) and size-exclusion chromatography, and was concentrated to 600 µM. UBE2T^C86K^ was conjugated to ubiquitin through an isopeptide bond in a 200 µL reaction containing 125 µM His-Ubiquitin (Boston Biochem U-530), 125 µM UBE2T^C86K^, 0.85 µM UBE1 (Boston Biochem E-304-050), 5 mM adenosine triphosphate, 5 mM MgCl_2_ in 50 mM CAPS, 150 mM NaCl, 0.2 mM TCEP, pH 10.0. After 4 hr incubation at 25 °C, the reaction was diluted to 1 ml in 50 mM Bicine, 150 mM NaCl, pH 8.0. The ubiquitin-conjugated UBE2T^C86K^ was separated from the non-ubiquitinated UBE2T^C86K^ by Ni-NTA chromatography, eluted by 20 mM Bicine, 150 mM NaCl, 200 mM imidazole, pH 8.0, then separated from UBE1 and ubiquitin by gel filtration.

### In vitro ubiquitination

Unless stated otherwise, mono-ubiquitination reactions contained 0.1 µM recombinant human Core complex, 1 µM substrate (ID, ID^-/Ub^, I^mut^D, FANCI or FANCD2 as indicated), 26 µM Ubiquitin (Boston Biochem U-530), 0.3 µM UBE1 (Boston Biochem E-304-050), 1.5 µM UBE2T (Boston Biochem E2-695), 2 µM dsDNA (58 bp), 5 mM adenosine triphosphate, 5 mM MgCl_2_ in 20 mM Tris-HCl, 100 mM NaCl, pH 8.0. 40 µL reactions were setup on ice and then incubated at 28 °C. At the indicated time points, a 3 µL aliquote was mixed with 9 µL NuPAGE LDS sample buffer (Invitrogen) and heated at 95 °C for 2 min to stop the reaction. Samples were separated by NuPAGE 3-8% Tris-Acetate SDS-PAGE (Invitrogen) in duplicate sets, and detected either with Coomassie blue staining (6.5 µL loading) or western-blotting (4.5 µL loading). Replacing the 58 bp dsDNA with 58 bp nicked dsDNA had no detectable effect on ubiquitination (not shown).

The ID^-/Ub^ substrate was prepared by first separating mono-ubiquitinated FANCD2^Ub^ from FANCI^Ub^ by anion exchange chromatography (MonoQ) of the ID^Ub^ complex, and concentrating it to 42 µM in 20 mM Tris, 350 mM NaCl, 10% glycerol, pH 8.0. FANCD2^Ub^ (3 µM final concentration) was then mixed with 6 µM dsDNA (58 bp), incubated for 10 min on ice before addition of a one molar equivalent of FANCI. After 30 min incubation on ice, the ID^-/Ub^ complex was mono-ubiquitinated as above at 1 µM concentration.

### Cryo-EM image processing

For all data sets, movie frames were aligned using MOTIONCOR2^32^, and the contrast transfer function parameters were estimated with CTFFIND4^33^. Initial 2D templates for autopicking were derived from manual picking of Core particles followed by 2D classification. For the Core complex, a total of 9760 aligned micrographs were used for autopicking using template matching in RELION3^34^, followed by reference free 2D classification. An initial 3D model was generated from 2D averages in EMAN2, followed by RELION3 auto-refinement. 3D classification in RELION3 was then used to remove particles in classes of poorly defined features or limited resolution that failed to significantly extend beyond the resolution at which the autopicking templates were low-passed. After applying Bayesian beam induced motion correction, scale and B-factors for radiation-damage weighting and per particle CTF refinement, an additional round of 2D classification was used to remove extant noise particles. The resulting data set of 671,927 particles was then used for a consensus reconstruction which resulted in a resolution of 3.4 Å (Extended Data Fig. 2a) were also applied before the consensus refinement which resulted a resolution of 3.4 Å. All reported map resolutions are from gold-standard refinement procedures with the FSC=0.143 criterion after post-processing by applying a soft mask.

At this stage, successive 3D classifications showed two kinds of conformational heterogeneity in the Core particle. One involved the inactive and active sides of the Core moving, mostly orthogonal to the flat shape (up and down in Figure 1a), relative to the central hub, and this was coupled to the movement of the two FANCA arms. The other involved the active-side interface between FANCA^CTD^’ and FANCG^CTD^’ (Extended Data 10h) dissociating, resulting in poor order and density for FANCA^CTD^’ and for the MD and CTD domains of the inactive side FANCA. Approximately 50 % of the particles contained an intact FANCA^CTD^’-FANCG^CTD^’ interface (FANCA conformation 1 in Extended Data Fig. 2a), and these were used for the focused refinement of FANCA. The remaining particles appeared to have a continuum of conformational states for FANCA^CTD^’-FANCA^MD^-FANCA^CTD^, and even after successive 3D classifications to identify a narrower range of conformational states (FANCA conformation 2 in Extended Data Fig. 2a), they could not be modeled reliably. The fraction of particles that adopt the FANCA-conformation 1 is comparable in the Core-alone, Core-ID and Core-UBE2T-ID data sets, and the functional significance, if any, of the FANCA^CTD^’-FANCG^CTD^’ interface slipping is unknown. The focused 3D refinements of the Core utilized seven non-overlapping soft-masks, except for the three FANCA-FANCA’ masks that overlapped partially (Core portions encompassed in masks and resulting FSC curves are shown in Extended Data Figs 2a and b, respectively). The 7,331 particles that correspond to a symmetric conformation of the Core (Extended Data Fig. 14) were identified using iterations of 3D classification, successively removing recognizable asymmetric Core particles. Their reconstruction was done using the RELION3 multi-body option with signal subtraction in point group C1, with three bodies corresponding to the left, middle and right side of Extended Data Fig. 14b.

The Core-ID and Core-ID-UBE2T data sets were processed similarly to the Core data, respectively yielding 289,005 and 295,721 clean particles from 4309 and 10932 micrographs after Bayesian polishing and CTF refinement. Because 2D and 3D classifications indicated that only a subset of particles contained bound ID, we performed partial signal subtraction of the consensus map outside the ID-FANCE^CTD^-FANCL’ or I ID-FANCE^CTD^-FANCL’-UBE2T density from each particle, followed by 3D classification without alignment to identify the ID-containing particles. We then performed focused 3D refinements on the original particles as with the Core data set, except for fewer masks (Extended Data Figs 2c and d).

### Model building and refinement

The Core structure was built into the maps with COOT^35^ and O^36^. We also utilized the available crystal structures of a FANCF C-terminal domain (residues 156-357, PDBID 2IQC), of fly FANCL (PDBID 3K1L) and human FANCL RWD2-RWD3 domains (also known as DRWD for Double RWD; PDBID 3ZQS). Initial structure refinement was done with REFMAC5^37^ modified for cryo-EM. For this, we first aligned the individual focused maps on the consensus reconstruction map in CCP4^38^, and then combined them with the composite sfcalc option of REFMAC5 to construct a single set of structure factors to 3.1 Å (Extended Data Fig. 2b, right). As the resolution of the FANCA-focused maps is lower (3.4, except for the FANCA^NTD^ which is to 3.7 Å), this results in higher temperature factors for the portions of the structure within these maps. Initial refinement with REFMAC5 in reciprocal space utilized NCS restraints for portions of subunits deemed to have the same conformation, and also secondary structure restraints generated by PROSMART^37^. Real space refinement was performed with the phenix.real_space_refine program of the PHENIX suite^39^. In the final stages the model was refined with REFMAC5, followed by TLS refinement. The disordered regions of Core subunits are indicated as dashed lines on Extended Data Figures 4a to h.

The structures of the Core-ID and Core-UBE2T-ID complexes were built using the refined Core structure, also utilizing the available crystal structures of human FANCL RING-UBE2T (PDBID 4CCG) and of the FANCE^CTD^ (PDBID 2ILR), as well as the cryo-EM structure of the human ID complex (ref). The Core subunits, the larger ones divided into individual domains, were fitted into the maps with CHIMERA, with the junctions between domains rebuilt as necessary. The models were refined in both reciprocal and real space as with the Core complex, except external structure restraints using the Core, ID, FANCE^CTD^ and UBE2T structures were also used because of the lower resolution of these maps.

### Data availability

The Core, Core-ID, Core-UBE2T-ID coordinates and the corresponding cryo-EM maps, including the focused reconstructions and the composite maps used in refinement, are being deposited with the Protein Data Bank and the Electron Microscopy Data Bank, respectively.

